# Tresor: An integrated platform for simulating transcriptomic reads with realistic PCR error representation across various RNA sequencing technologies

**DOI:** 10.1101/2025.03.15.643015

**Authors:** Jianfeng Sun, Adam P. Cribbs

## Abstract

The rapid advancement of high-throughput sequencing technologies has spurred the development of numerous computational tools designed to identify gene expression patterns from growing datasets at both bulk and single-cell sequencing levels. The recent advent of longread sequencing technologies has further accelerated the availability and refinement of these tools. The lack of ground-truth labels and annotations in sequencing data presents a significant challenge for evaluating the efficacy of analytical tools. To address this, we developed Tresor, an integrated platform for simulating both short and long reads at bulk and single-cell levels. We devised a tree-based algorithm to significantly accelerate in silico experiments at high PCR cycles. Tresor allows for customising sequencing libraries with highly modular and flexible read structures, facilitating the verification of sequencing-related biological discoveries. This tool also includes features that introduce substitution, insertion, and deletion errors at various stages of library preparation, PCR amplification, and sequencing, enhancing its applicability in diverse experimental conditions and simulating real world conditions. Our results demonstrate that, upon removal of PCR duplicates, cell type-specific gene expression profiles derived from our simulated reads highly resemble reference data. We envisage that Tresor will provide valuable insights into a broader range of transcriptomics analyses and support the development of more effective algorithms for read alignment and UMI deduplication.

## Introduction

RNA sequencing (RNA-seq) has revolutionised the study of the transcriptome (Stark et al. 2019; Ozsolak and Milos 2011), laying the foundation for further elucidating the functional roles of sequenced components across a wide range of biological processes (Kukurba and Montgomery 2015; Khoogar et al. 2022; Klein et al. 2015). Over recent decades, advances in RNA-seq technologies have enhanced our ability to investigate genetic regulation at various resolutions, ranging from bulk tissue samples (Wang et al. 2009) to individual cells (Macosko et al. 2015), and with different read-lengths, including short-read (Logsdon et al. 2020) and long-read capacities (Philpott et al. 2021)(Li and Wang 2021; Hrdlickova et al. 2017). The introduction of high-throughput sequencing technologies has precipitated a substantial increase in the volume of sequencing data (Andrews et al. 2021a; Bansal and Boucher 2019), with a concomitant rise in the number of computational tools for analysing the data (Andrews et al. 2021b). Notably, the recent advent of long-read sequencing technologies has further accelerated the availability and refinement of these tools (Wang et al. 2021; Dong et al. 2023; Gamaarachchi et al. 2022). However, the absence of ground-truth labels and annotations in the sequencing data presents a major challenge in evaluating the performance of these tools (Lähnemann et al. 2020). This lack of standardisation hampers the development of universally accepted analytical approaches. Consequently, there is a critical need for well-annotated datasets to accurately evaluate and improve sequencing methodologies.

The use of simulated sequencing data has become a standard approach for assessing the reliability of transcriptomics analysis methods (Smith et al. 2017; Sun et al. 2024b). However, a range of available simulation methods and tools remains limited (Yu et al. 2020; Yan et al. 2023), with many tailored to domain specific tasks such as bulk long-read RNA-seq (Karaoğlanoğlu et al. 2024), short-read droplet-based single-cell RNA-seq (scRNA-seq) (Sarkar et al. 2019), or bulk short-read transposase-accessible chromatin using sequencing (scATAC-seq) (Chen et al. 2021). The realistic demands for testing transcriptomics analysis methods and supporting customised sequencing experiments far exceed these available simulation tools (Cao et al. 2024; Simmons et al. 2023; De Rop et al. 2023). These experiments require flexible modification in sequencing protocols across various technologies (Philpott et al. 2021; Karst et al. 2021; Picelli et al. 2014; Lu et al. 2011; Zhang et al. 2017; Choi et al. 2022). For example, our recent study demonstrates that bespoke *in silico* simulations can determine optimal positions for anchor sequences in oligonucleotide synthesis to mitigate errors (Sun et al. 2024a). Additionally, these bespoke *in silico* simulations are supportive of quantifying the contribution of bead synthesis errors to the overall sequencing error profile. Based on the fact that one out of ten Drop-seq beads suffering from deletion errors according to (Zhang et al. 2019), our simulations initiated with 50 molecules at one gene-by-cell type estimate around 8-14% of final sequencing reads suffering from bead synthesis-induced errors, and notably, this figure can even become doubled if the PCR error rate is elevated by one order of magnitude (i.e., 1e-4 to 1e-3). Nonetheless, current tools gain insufficient momentum for their capability to customise sequencing libraries across, and, more importantly, no tools are dedicated to simulating errors (insertions, deletions, and substitutions) in complex conditions (Choi et al. 2022).

Most available tools fail to simulate reads that have undergone multiple PCR amplification cycles. The primary constraint is the exponential increase in time complexity associated with simulating reads across successive PCR cycles. This problem intensifies as additional PCR cycles are required to generate the necessary amplified input. Addressing this limitation is crucial for improving the accuracy and reliability of transcriptomics analysis methods. In certain sequencing applications, such as single-cell RNA sequencing (scRNA-seq) at the long-read level, high PCR cycles are essential for amplifying RNA from a single-cell to a measurable level. (Rodger et al. 2024). For instance, PCR-cDNA sequencing of Oxford Nanopore sequencing technologies (ONT) relies on extensive PCR amplification (Grünberger et al. 2022; Bayega et al. 2022). However, PCR amplification can introduce errors into reads (Marx 2017; Islam et al. 2014; Michlits et al. 2017; Zurek et al. 2020), and the extent to which these errors propagate through successive PCR cycles is not well understood.

Our recent study utilises homotrimer unique molecular identifiers (UMIs) and common molecular identifiers (CMIs) to reveal that PCR introduces 15-40% inaccuracies, predominantly affecting molecular counting (Sun et al. 2024b). This work demonstrates that the use of homotrimer UMIs effectively removes PCR artefacts. Despite these advancements, current simulation tools remain inadequate in addressing these fundamental questions, underscoring the need for improved methods to simulate the impact of multiple PCR cycles on sequencing accuracy.

In this study, we present **T**ree-based **re**ad **s**imulat**or** (Tresor) that employs a tree structure approach to model the trajectories of sequencing reads across PCR cycles, thereby facilitating faster and more efficient simulation of sequencing reads. This tool proposes that the PCR-amplification of sequencing molecules can be effectively represented using a tree-based data structure. By iteratively traversing the leaves of these trees with their depths,

Tresor optimises both of simulation speed and the consumption of computational resources. Our results show that tree-based methods significantly speed up read simulation compared to non-tree based approaches, particularly when simulating reads across a high number of PCR cycles. Tresor is destined diverse sequencing applications, applicable to both single-cell and bulk sequencing at various levels, including short-read and long-read RNA-seq technologies. Moreover, it can simulate multisource errors that are introduced in library preparation, PCR amplification, and sequencing, providing rough estimates for customised sequencing experimental setups. This work can enhance the reliability and robustness of methods aimed at correcting sequencing errors, thereby enabling more accurate interpretation of experimental results and facilitating advancements in molecular biology research.

## Results

### Tresor Implementation

Tresor is designed to simulate either short-reads or long-reads under bulk RNA-seq and scRNA-seq conditions across six scenarios: specifically by varying sequencing errors, PCR errors, lengths of unique molecular identifiers (UMIs), sequencing depths, numbers of PCR cycles, and PCR amplification rates, respectively (**Fig. 1**). Tresor is implemented with Python, which can be accessed through both Python and Shell commands. The software outputs sequencing libraries that include comprehensive annotations, enabling the tracking of both the original and PCR-amplified molecules under complex conditions, such as sample multiplexing with Unique Molecular Identifiers (UMIs) and cell barcodes. These features are instrumental in quantifying changes in the number of PCR duplicates across different experiment settings. By providing these detailed annotations, Tensor supports both the evaluation of existing UMI deduplication tools and the spur of the development of novel methodologies.

**Figure 1.**
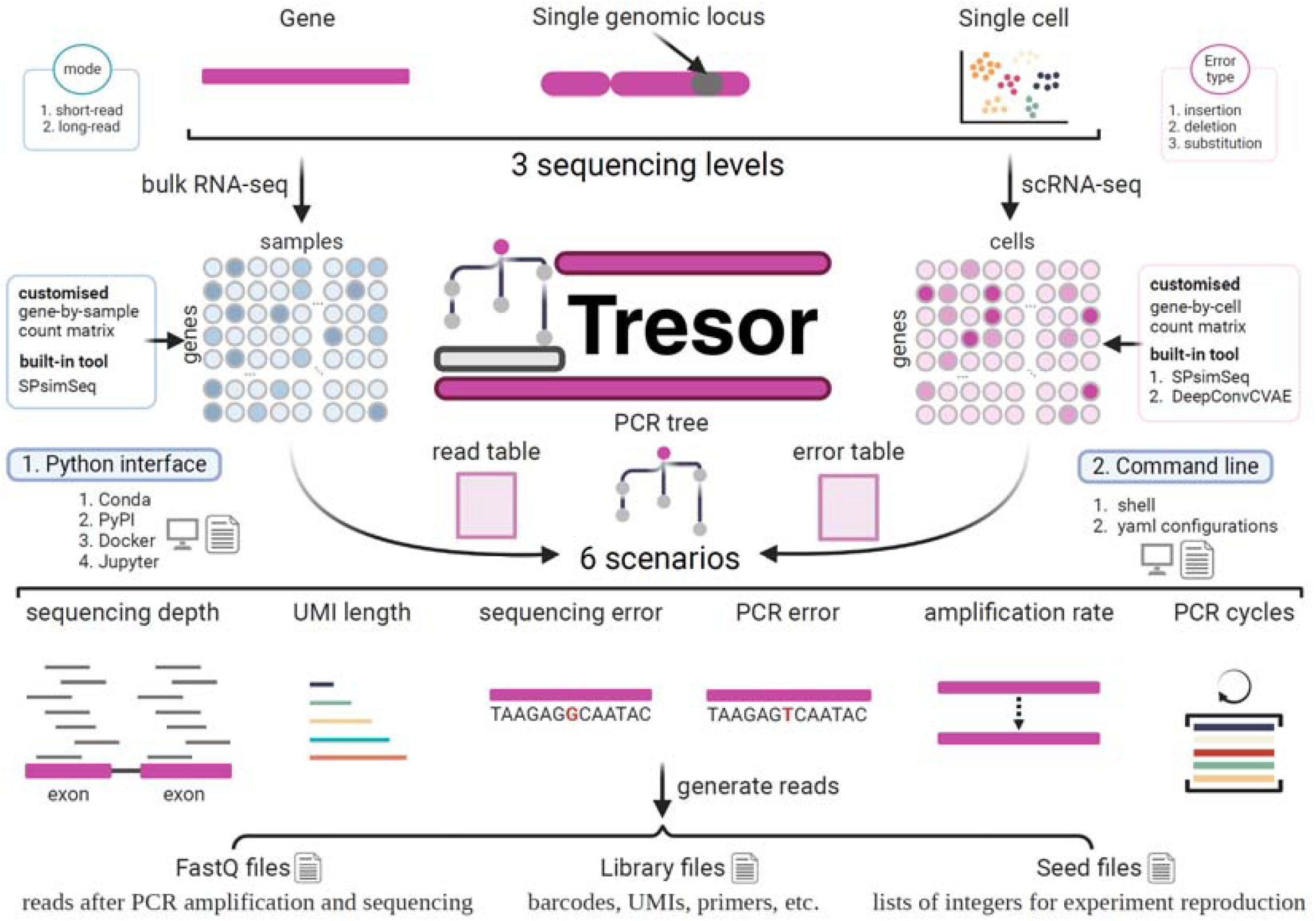
Tresor enables read simulation with ground truth at multiple sequencing levels in various sequencing scenarios.

In particular, to simulate reads in multiplex sequencing, Tresor uses an external tool SPsimSeq to generate cell-by-gene count matrix and sample-by-gene count matrix. At the single-cell level, Tresor generates cell barcodes that are used to tag reads in two ways, that is, randomly synthesising a barcode for each cell or selecting a barcode in line with 10X Genomics (V2 or V3 chemistry) (Vandereyken et al. 2023) and Dropseq protocols

(Bageritz Josephineand Raddi 2019)

. At the bulk level, Tresor assigns a unique mark to each read from the same sample rather than automating the attachment of molecular barcodes to reads.

Tresor is designed as a modular tool, comprising three distinct components: simulation of sequencing libraries, generation of count matrices, and amplification and sequencing of reads. Configuration of most parameters related to PCR amplification and sequencing is managed through a YAML file, allowing flexible adaptation to different experimental setups. For preparing a library tailored to bulk RNA-seq, users can employ the *library_bulk* module as demonstrated below. This module architecture not only enhances usability but also facilitates customisation for specific sequencing applications:

*tresor library_bulk -cfpn library*.*yml -snum 50 -nspl 10 -ngene 10 -gsimulator spsimseq -rfpn R/R-4*.*3*.*2/ -permut 0 -sthres 3 -wd ./tresor/data/simu/ -md short_read* where *cfpn* represents the location of configuration file, *snum* represents the number of initial molecules to be sequenced, *nspl* represents the number of samples, *ngene* represents the number of genes, *gsimulator/rfpn* represents the name/location of a tool for count matrix generation, *permut* represents permutation times, *sthres* represents the threshold to generate UMIs between a certain Hamming distance, *wd* represents a working directory, and *md* represents whether to generate short reads or long reads. The resulting files contains the libraries of different read components (e.g., UMIs and barcodes) and the seeds. Upon generation of the above sequencing library, then, as an example to generate FastQ reads across various sequencing errors, we can utilise the *seqerr_gene* module as follows.

*tresor seqerr_gene -cfpn bulk*.*yml -snum 50 -permut 0 -sthres 3 -wd ./tresor/data/simu/ -md short_read*

### Read simulation methods

Read simulation, especially at elevated PCR cycles, demands considerable computational resources in terms of CPU and memory. To reduce the computational costs, we devise a range of read simulation methods that vary in their strategies for efficiently amplifying and storing reads, as well as generating and assigning errors. These methods leverage various strategies including the construction of PCR trees, read tables, and error tables, which facilitate the accurate propagation of read and error data through successive PCR cycles. Consequently, we introduce three distinct approaches: PCR tree-based, read table-based, and error table-based methods, each tailored to reduce CPU and memory demands while improving simulation fidelity.

#### PCR Tree-based methods

Tree-based data structure has been widely used to address many biological problems, such as efficient searching operations (Emms and Kelly 2022; Ondov et al. 2016; Varón and Wheeler 2013) and hierarchical representation of information (Gundem et al. 2015; Jahn et al. 2016; Huang et al. 2021). In this context, PCR amplification is analogous to tree structures where exponential duplication of molecules corresponds to the branching of a tree. Each PCR cycle can be represented as a layer in the tree, with amplified reads depicted as nodes. Following PCR amplification, partial reads will randomly be subsampled for sequencing as in (Smith et al. 2017; Best et al. 2015; Sarkar et al. 2019). Thus, the primary challenge in this method arises from the inefficient use of computational resources when assigning errors to reads that are not ultimately used for sequencing. To mitigate this, our approach avoids the generation and assignment of errors during the PCR amplification phase. Instead, this process only records the PCR cycle numbers required for a molecule to amplify itself, thus streamlining the highly laborious process of read replication inherent to PCR amplification. Specifically, a read selected for sequencing ends up with a string of dash-delimited digits (e.g., 1-2-5-6-8) where each represents a PCR cycle that an ancestor of the read goes through for its replication.

A further challenge is ensuring that molecules amplified in any given PCR cycle *i* originate strictly from PCR cycle *i*-1, as errors from all the previous cycles are inherent in these molecules. To address the lack of effective strategies for tracing read information when errors are not assigned during amplification, we employ noel methodology. We generate a unique tree for each molecule sampled for sequencing, storing all intermediate reads. This allows for tracing different versions of an initial molecule through various PCR cycles (**Fig. 2a)**. In this PCR tree, different versions of a molecule are connected by unique and shared paths, enabling the tracing of read information through specific paths.

**Figure 2.**
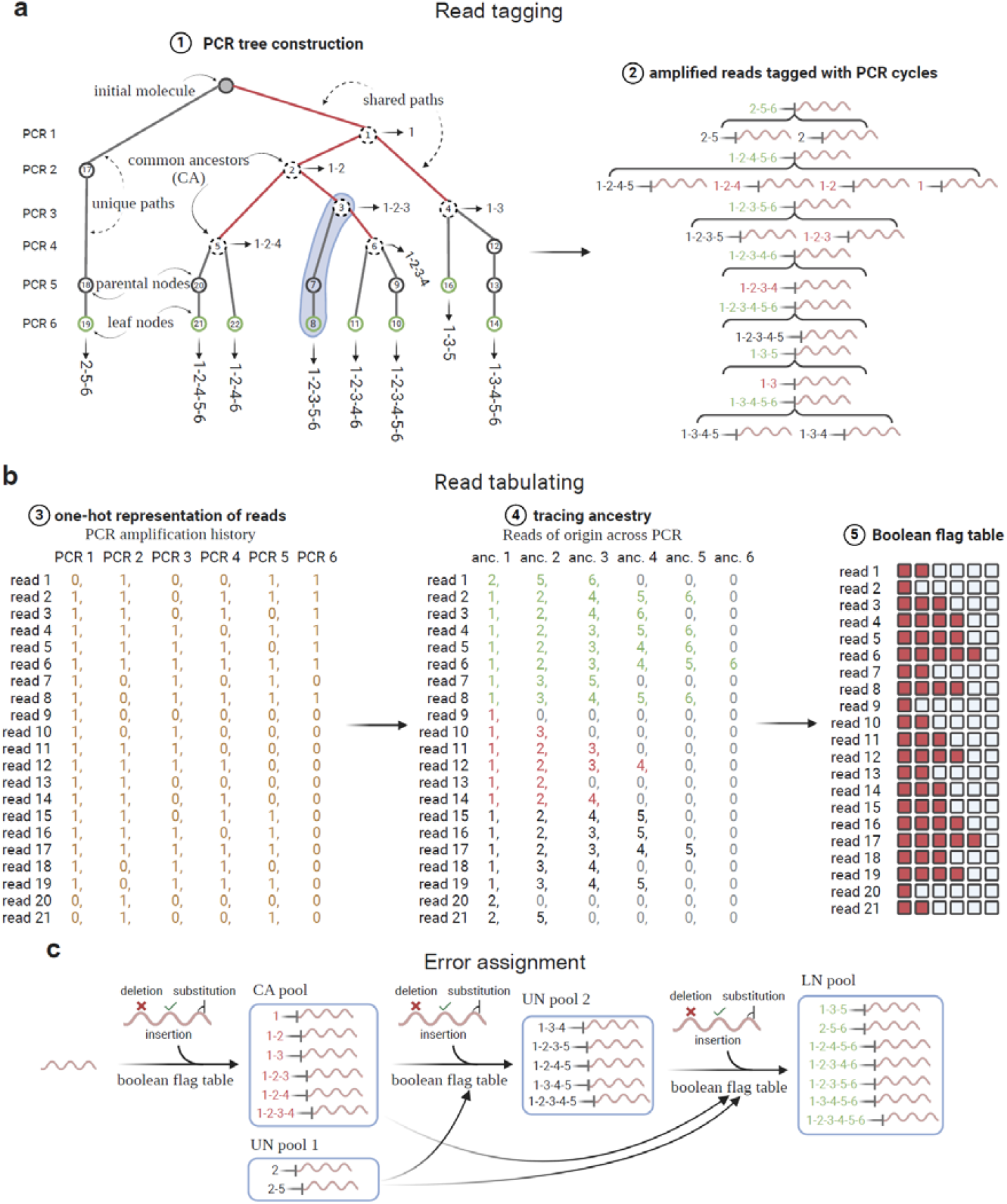
Illustration of the *bfTree* method for read simulation based on the PCR tree. **a.** PCR tree construction. Reads are tagged with a concatenation of identifiers of PCR cycles (separated by underscore) through which they are PCR amplified. The reads circled in green are those present in the final sequencing pool. By contrast, the rest of reads located at the parental nodes are intermediate versions of the initial molecule, which we present to illustrate how the final sequencing reads in green use them as proxies to trace the correct ancestors across the PCR tree. **b**. Tabulating reads and constructing the Boolean flag table for tracing the ancestry of reads. **c**. schematic workflow for assigning errors to reads based on the ancestral relationships between reads in the PCR tree. CA: common ancestor. UN: unique node. LN: leaf node. Blocks in red in the Boolean flag table represents True and False, otherwise. For instance, read 4 (with a PCR amplification history of 1-3) will be used for generating its following 4 reads without visiting other versions of the molecule. In addition, the highlighted region demonstrates that read 4 is used twice for generating read 12 and read 16. The right panel of a shows exactly the least number of steps required to amplify all reads located at leaves of the tree.

To facilitate this tracing and manage the complexity, we have developed two specific methods:

1. *bfTree*. This method uses a Boolean flag table (*bfTree*) where each column corresponds to a PCR cycle. Boolean flags indicate whether reads have undergone amplification in identical preceding PCR cycles. This method ensures that errors, once generated, are assigned consistently to the correct target read, preventing misallocation of errors across different read versions. This technique is detailed in **Fig. 2 and Supplementary Table 1**.
2. *spTree*. To improve efficiency, particularly when dealing with large numbers of reads related to the same molecule, we developed *spTree*. This approach utilises a read cache table to lazily calculate a short path for error assignment to reads (see **Supplementary Table 2**). This method avoids the time-intensive construction of Boolean flags, instead querying the cache table to ensure that errors are assigned using the same read version through successive PCR cycles (**Fig. 3a**).

**Figure 3.**
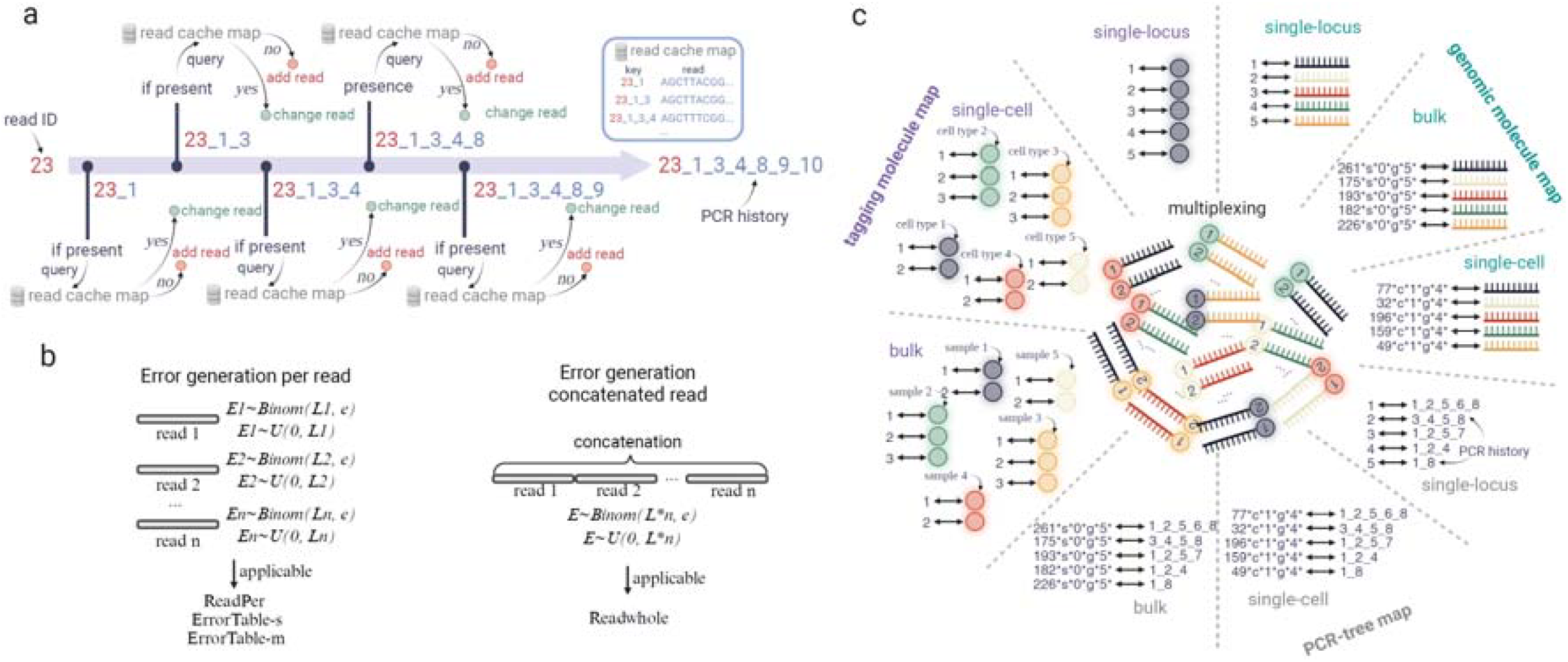
Profiles of sample multiplexing. a. Illustration of the *spTree* method for read simulation based on a read cache map. b. Strategies for error generation. Error generation per reads applies to the *ReadPer, ErrorTable-s*, and *ErrorTable-m* methods. c. Sample multiplexing. Genomic sequences are attached with UMIs and cell/sample barcodes. In Tresor, a read is tagged with a string that contains information regarding the types and identifiers of the sample, cell, and gene of origin.

For comparison purposes, we developed another two categories of methods, read table-based methods and error table-based methods, which differ in whether or not to assign errors during PCR amplification.

#### Read table-based methods

Read table-based methods (*preEAssign*) generate and assign errors during PCR amplification, which contain two strategies *ReadPer* and *ReadWhole*. As shown in **Fig. 3b**, the *ReadPer* strategy samples errors per read (see **Supplementary Table 3**), while the *ReadWhole* strategy samples errors for the concatenated sequence of all reads at once (see **Supplementary Table 4**). To achieve this, all necessary materials, including the sequences and identifiers of all reads, are passed onto the amplification stage, which depletes plenty of memories.

#### Error table-based methods

In contrast, error table-based methods (such as *postEAssign*) are memory-efficient since they do not require read sequences as input, and error assignment is deferred until the sequencing stage. Instead, these methods focus on documenting errors in error tables, which are then utilized during sequencing. This category includes two methods: *ErrorTable-s* and *ErrorTable-m* (see **Supplementary Table 5**). The former method removes from the error tables those reads that do not contain any errors, thus conserving memory. Conversely, the latter method retains all information about the reads in the error tables, without discarding any data.

### Runtime of read simulation methods across PCR cycles

We compared the runtime of read simulation by varying PCR cycles from 6 to 24 according to our specific simulation techniques used. To this end, we first generated a sequencing library that contains 50 sequencing molecules observed at a single genomic locus. We then recorded the runtime at each of those predefined cycles through which reads in the sequencing library are PCR amplified (**Fig. 4**). With the increase of PCR cycles, the exponential growth of runtime for all methods have been proven. We show that two tree-based methods significantly outperform both *preEAssign* (*ReadWhole* and *ReadPer*) and *postEAssign* (*ErrorTable-s* and *ErrorTable-m*) methods. Particularly, after 15 cycles, the tree-based methods can bring down the runtime by nearly two orders of magnitude as other methods can.

**Figure 4.**
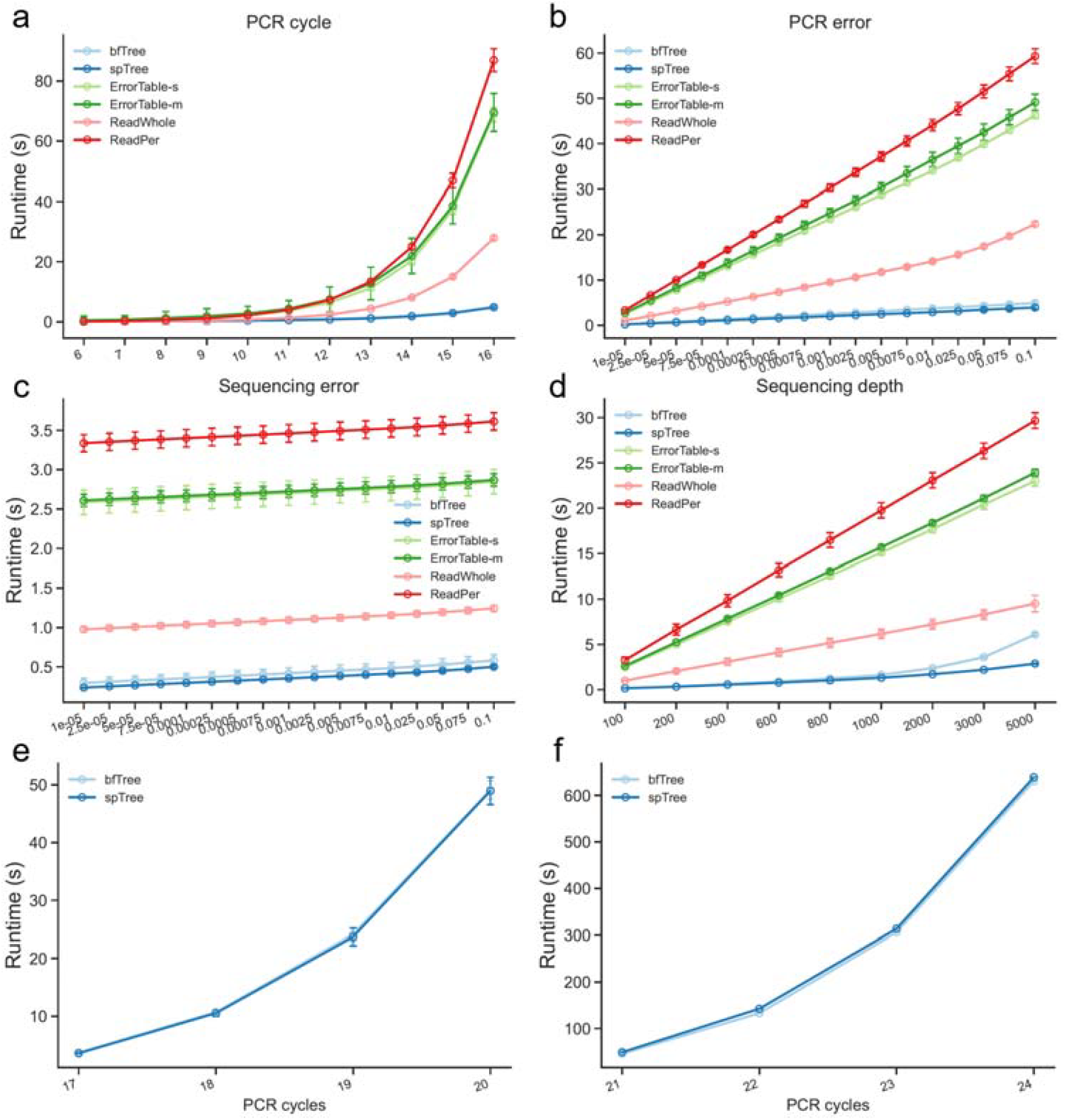
Runtime of different simulation methods at varying PCR cycles (a), PCR errors (b), sequencing errors (c), and sequencing depths (d). Runtime of bfTree and spTree at 17-20 PCR cycles (e) and 21-24 PCR cycles (f).

Moreover, non-tree-based methods exhibited significant challenges in amplifying reads beyond 16 cycles, particularly when the number of reads exceeds 1 million, with an amplification efficiency rate of 0.85. In contrast, tree-based methods demonstrated the capacity to efficiently generate sequenced reads from 1 million (16th cycle) to 10 million (20th cycle) PCR amplified reads without difficulty (**Fig. 4e**). The *preEAssign* and *postEAssign* categories of read simulation methods were excluded from the comparison due to extremely high runtime. Our results show that at 24th PCR cycle, tree-based methods can generate sequenced reads from around 0.13 billion PCR amplified reads in around 600 seconds (**Fig. 4f**), significantly accelerating read simulation. Notably, due to the elimination of read sequences during PCR amplification, tree-based methods exhibit minimal memory usage, even when processing 0.13 billion reads. Overall, our simulation demonstrates the effectiveness of Tresor in addressing the challenges associated with the exponential growth in time complexity of PCR amplification. This makes it a valuable tool for preliminary computational analysis and verification of sequencing experiments that require high PCR cycles, such as deep scRNA-seq.

Moreover, it was found strenuous for non-tree-based methods to amplify reads starting from 16 cycles as the number of reads exceeds 1 million with an amplification efficiency rate of 0.85. By contrast, the tree-based methods were shown effortless to generate sequenced reads from 1 million (16th cycle) to 10 million (20th cycle) PCR amplified reads (**Fig. 4e**). **The *preEAssign* and *postEAssign* categories of read simulation methods are opted out of the comparison due to extremely high runtime**. Our results show that at 24th PCR cycle, the tree-based methods are able to generate sequenced reads from around 0.13 billion PCR amplified reads in ∼600 seconds (**Fig. 4f**), significantly speeding up read simulation. Importantly, the tree-based methods have minimal memory usage as we avoid using read sequences during PCR amplification, even when dealing with 0.13 billion reads. Altogether, our simulation suggests the effectiveness of Tresor in wrestling with problems brought on by the exponential growth nature of the time complexity of PCR amplification, promising to be supportive of the preliminary computational analysis and verification of sequencing experiments calling for high PCR cycles, such as deep scRNA-seq.

### Analysis of computational costs of read simulation methods

As expected, we discovered that introducing errors during PCR amplification constitutes the most time-intensive step in the process of read generation across all application scenarios (**Fig. 4a-d**). These methods belong to the *preEAssign* and *postEAssign* categories, which draw error positions and error nucleotide types from statistical distributions during PCR amplification no matter whether they assign the error nucleotide types to the error positions before or after PCR amplification. This observation indicates that time optimisation can be made by altering when to assign errors since molecules in the final sequencing pool are underrepresented compared to those from PCR amplification, leading to a resource-saving strategy. By comparing the *ReadWhole* method with *ErrorTable-s, ErrorTable-m*, and *ReadPer*, we found that the time required for simulating errors according to per read is significantly higher than that according to the concatenated sequence of all reads, especially when the conditions become more restricted (e.g., high PCR errors) in most of the scenarios. This arises from the vastly reduced operations on generating distributions to simulate errors. While merging reads as a whole using ReadWhole incurs additional resource consumption, the subsequent single-error generation dramatically reduces the overall time cost compared to those methods drawing errors multiple times.

### Variation tendency for runtime

We investigated the variation in runtime for read simulation across different parameters and application scenarios (**Fig. 4**). Our analysis revealed that the runtime of most methods increases linearly with the rise in PCR errors, sequencing errors, and sequencing depth, but exponentially with the number of PCR cycles. The runtime relatively insensitive to changes in sequencing errors, likely due to the selection of a set of underrepresented reads for sequencing in the final pool, thus stabilising the overall computational cost. Additionally, we observed that as sequencing depth increases, particularly beyond 1000, the runtime of the *bfTree* method exhibits a noticeable nonlinear variation compared to the *spTree* method. We systematically benchmarked the runtime of simulating reads across a range of amplification efficiencies from 0.1 to 1. The problem becomes more pronounced when a larger number of reads are selected for sequencing, attributed to the sharply increasing cost of generating the Boolean table to track their usage and status across the PCR tree.

### Examination of read simulation efficiency for multiplex sequencing

We subsequently benchmarked the runtime for simulating reads at both the bulk and single-cell sequencing levels. An initial count matrix for bulk RNA-seq was simulated with 10 samples and 10 genes using the *SPsimSeq* tool. Similarly, a count matrix for scRNA-seq with 10 cells and 10 genes was generated. Our data shows that, using an initial pool of 34,496 expression counts as input, methods from the *preEAssign* and *postEAssign* categories can generate the reads using 6-7 PCR cycles within a few seconds at the bulk level. Notably, tree-based methods achieve this within one second (**Fig. 5a**). At the single-cell level, tree methods also exhibited rapid read generation, with times ranging from 1s (6 PCR cycles) to 18 (12 PCR cycles) seconds (**Fig. 5b**). Additionally, all methods adhere to the exponential growth rule regarding the time complexity, consistent with observations at a single given genomic locus. Specifically, the average runtime of tree-based methods is approximately 8.1 millisecond per non-zero sample-by-gene position at the bulk level, and about 7.5 (6 PCR cycles) millisecond and 0.44 (12 PCR cycles) second at the single-cell level for each of 40 cells where at least one gene is expressed. These results suggest that tree-based methods are highly efficient for data generation requiring extensive PCR-cycle simulation.

**Figure 5.**
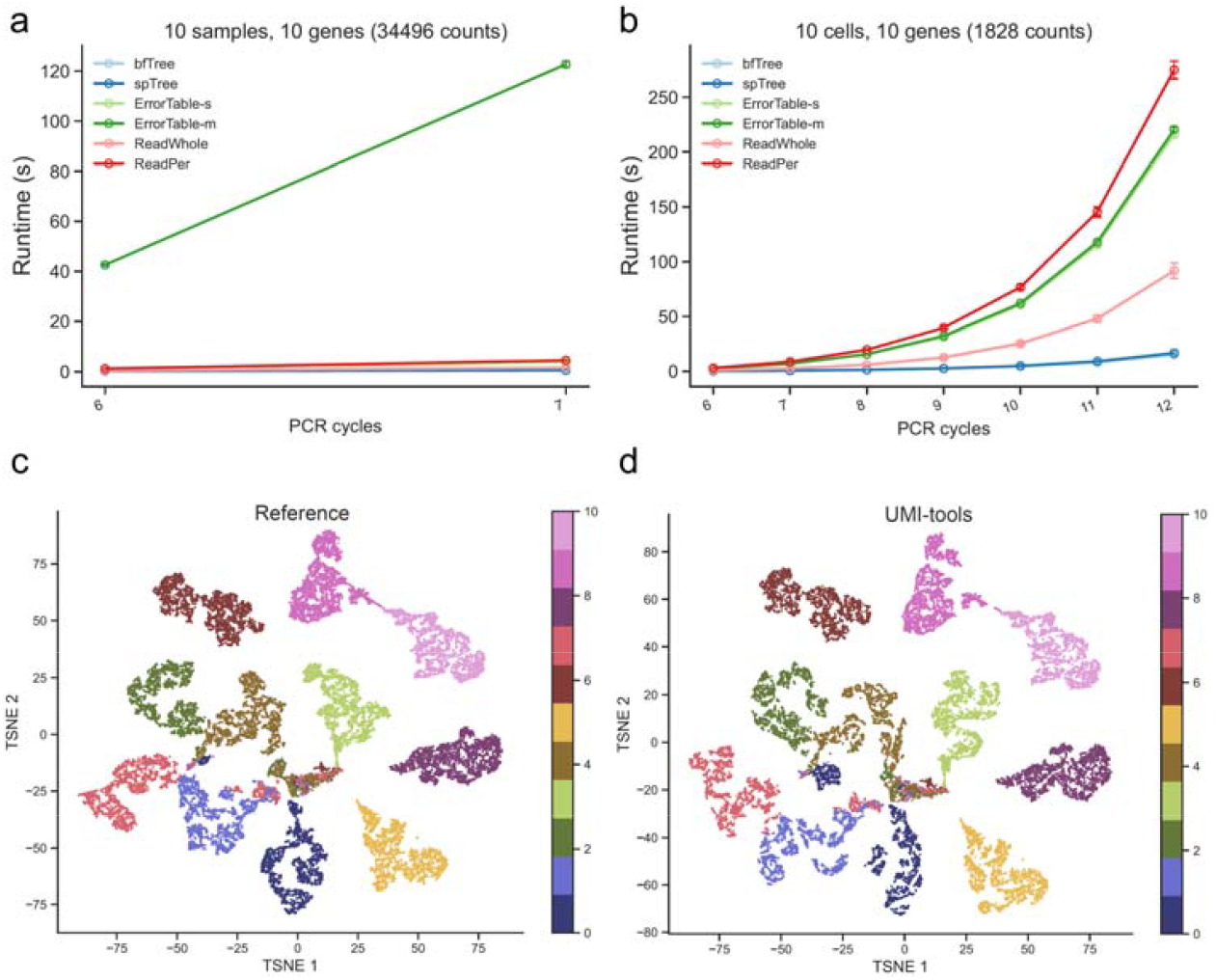
Analysis of simulating reads at the bulk and single-cell levels. Runtime of simulation methods at varying PCR cycles using bulk RNA-seq data (a) and scRNA-seq data (b). c. UMAP plot of reference scRNA-seq data. d. UMAP plot of simulated sequencing data after removing PCR duplicates with UMI-tools.

### Assessment of read quality at the scRNA-seq level

To understand whether simulated reads through our constructed sequencing simulation framework make biological sense, we evaluated the effect of removing PCR duplicates from the simulated reads and examined it at the single-cell level due to the ease of visualisation. To compare with the deduplicated counts, we first generated a count matrix as ground truth using our in-house simulation tool *DeepConvCVAE*, which annotates each cell with its corresponding cell type, as shown in **Fig. 5c**. This count matrix is constructed with 22,000 cells (11 cell types, 2000 cells per cell type) and 20,736 genes. Then, the gene expression value (initial UMI count) at each cell-by-gene position took turn being subjected to library generation, PCR amplification, and sequencing, as used in Tresor. For convenience, the resulting FastQ reads simulated at a cell-by-gene position were repeatedly used for other positions that accommodate a gene expression value that is identical to the one in that position. Finally, we applied the *Directional* methods in UMI-tools to remove PCR duplicates from the simulated reads to obtain the ultimate molecular counts. The data was projected onto a UMAP. As can be seen in **Fig. 5d**, the fidelity of molecular quantification on the simulated data is well-preserved, suggesting that our Tresor framework is capable of supplying biologically meaningful data and the simulated reads can, to a large extent, be put into practice in sequencing-involved problems.

## Methods

### Mathematical framework of read simulation

Stochastic modelling of branching processes has been extensively employed to capture the inherent variability in the reproduction and growth of populations (Lalam 2006). PCR amplification or the accumulation of mutations resembles a branching process, as the replication of reads or the occurrence of errors arises from randomness (Sun 1995). Therefore, we estimated a few key parameters, such as the quantities of reads and mutations, from stochastic models of the branching processes.

#### Read amplification

Let *N*_*i*_ be the number of reads at PCR *i*. To quantify the total number of reads *N*_*i*+1_ at the next PCR cycle, the following branching process is introduced.

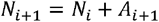

where *A*_*i*+1_ is the number of amplified reads at PCR cycle *i* +1 to be chosen from *N*_*i*_ using a constant PCR efficiency *a*. It has been widely suggested that the number of reads to be and Zador 2015). Hence, *A*_*i* +1_ can be binomially drawn by amplified is subjected to binomial distributions (Lalam 2007; Wang et al. 2000; Kebschull

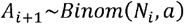

Then, we use a discreate uniform distribution to determine specific identifiers of *A*_*i* +1_ reads for amplification, which is expressed as

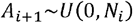

Alternatively, we implemented a uniformly sampling strategy in Tresor to determine the reads for amplification. The above branching process is rewritten to be

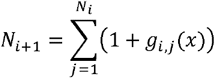

Where *g*_*i,j*_ (*x*) is an *indicator function*, which can only take two values: 1 if a read *j* at PCR cycle *i* is chosen for amplification at PCR cycle *i* +1 and 0, otherwise, such that

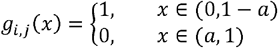

To solve *g*_*i,j*_ (*x*), we generate *N*_*i*_ probabilities for the random variable *x* by using a continuous uniform distribution as follows.

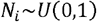

The working principle is that each read will be asked about whether it is selected for amplification: 1 when its assigned probability falls within the (0,1 − *a*) interval and 0, otherwise. Due to the possible deviation of the estimated number of reads from a binomial distribution (WEISS and VON HAESELER 1995), this approach is not typically favoured as the default choice for read amplification in Tresor.

#### DNA polymerase errors

Let *M*_*i*_ be the total number of DNA polymerase errors stochastically occurring at PCR *i*. Likewise, the aggregation of errors at PCR cycle *i* +1 of is described as

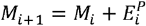

where 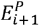 is the number of DNA polymerase errors that newly occurs in *A*_*i*_ amplified reads at PCR cycle *i* +1. Suppose that the sum of sequence positions of *A*_*i*_ reads is 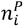 and the constant DNA polymerase error is 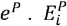 has widely been reported to be binomially distributed (Orton et al. 2015; Drummond et al. 2005; Kou Ruqin AND Lam 2016), such that,

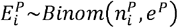

Besides, in another research (Rabadan et al. 2018), negative binomial distributions are suggested to be used for error modelling. To offer versatility, both methods are implemented as sampling error counts in Tresor. The distribution is often employed to quantify the occurrences of failures from a series of independent and identically distributed Bernoulli trials until a number of successes are achieved. Hence, we calculated the number of successfully synthesised nucleotides at PCR cycle *i* +1 by 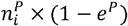,denoted as 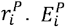 is drawn from the following distribution.

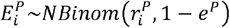

Then, we use a discreate uniform distribution to determine 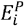 sequence positions from 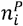, which is expressed as

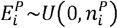

#### Sequencing errors

Following PCR amplification, we performed subsampling of reads from PCR product for sequencing, as with this study (Best et al. 2015). Let *B*_*i*_ be the total number of subsampled During sequencing, *E*^*s*^ errors are introduced and modelled using a binomial distribution.

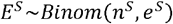

where *e*^*s*^ is the sequencing error rate and *n*^*s*^ is the number of nucleotides of *B*_*i*_ reads. Then, we use a discreate uniform distribution to determine *E*^*s*^ sequence positions where errors occur in their nucleotides, which is written by

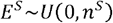

### Library preparation and generation

We provided a diverse range of functionalities for library preparation and generation to better cater to the requirements of data simulation across various sequencing technologies and experimental settings. To simulate long reads and short reads, we offered two modules: one for extracting sequences from a reference genome and another for randomly synthesising sequences (**Fig. 6**). A reference genome can stem from any species and is required to be organised in the Fasta form. Randomly synthesising sequences will require sampling of nucleotides from a uniform distribution.

**Figure 6.**
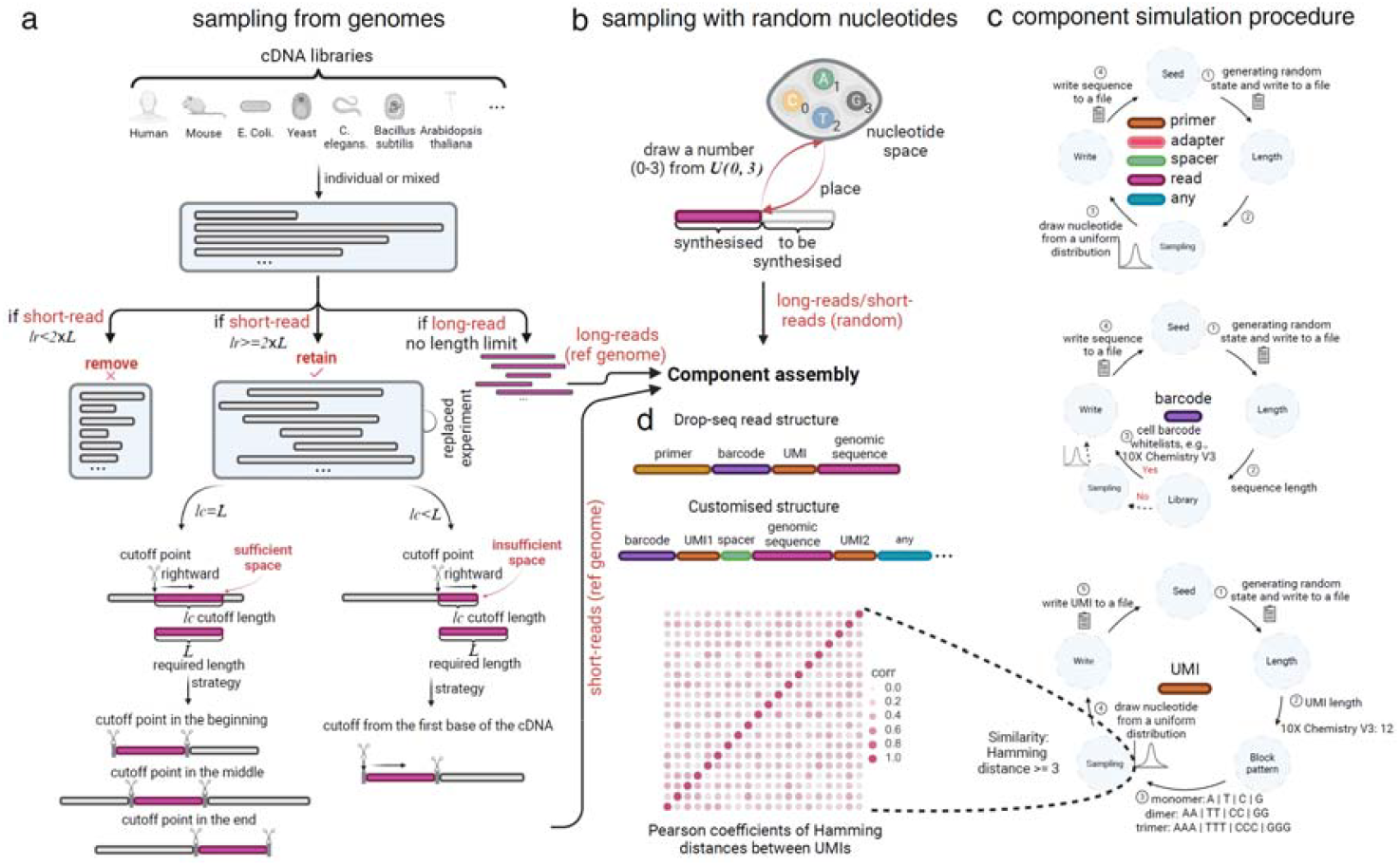
Workflow for library simulation. a. Sampling the full-length or partial cDNAs from a reference genome for long-reads or short-reads, respectively. b. Randomly synthesising full-length or partial nucleotide sequences for long-reads or short-reads, respectively. c. Procedures for simulating read components, such as primer, adapter, spacer, and UMI. d. Assembly of various read components.

#### Long-reads

If the working mode is set to generating long-reads with a reference genome supplied, Tresor will first build an index to enable a fast search for genomic sequences with their names. If without a reference genome supplied, Tresor will randomly synthesizing length-predetermined long-reads.

#### Short-reads

Similarly, Tresor sets out to generate short-reads under two working modes: with or without a reference genome. In the absence of a reference genome, Tresor synthesizes sequences randomly by sampling nucleotides of a fixed length. This length is set to be compatible with Illumina sequencing by default. If a reference genome is specified, a sampling strategy is applied to pick out fixed-length sequences from reference sequences. Tresor will exclude reference sequences that fall below a specified length threshold to avoid potentially prolonged processing times when acquiring short sequences with appropriate cutoff points. The length threshold is set to the doubled short-read length. When the cutoff point, which lies somewhere within the raw reference sequence, cannot dispose of the extraction of fixed-length short sequences, we employ a strategy where nucleotides are counted from the beginning of the sequence until the desired length is reached for the short sequence.

Outside of simulating genomic sequences, Tresor provides a variety of commonly used read components designed for capturing molecules of interest or facilitating partitioning between adjacent components. These include adapters, UMIs, barcodes, and more. (**Fig. 6c**). The majority of them are synthesized using the same procedures, encompassing seeding, length checking, nucleotide sampling, and file writing. However, owing to their distinct functions, a minority of them (e.g., UMIs) are generated with additional constraints, such as similarity thresholds. To make simulation experiments reproducible, Tresor saves all seeds used for generating all of the read components.

#### UMIs

UMIs are indicative of the identities of their attached molecules. Established research highlights their importance in eliminating PCR duplicates to ensure accurate molecular counting. However, UMIs simulated during library preparation may exhibit high similarity, which are exacerbated by the error-prone read generation process. To enhance their distinctiveness, we impose restrictions on their similarity (see **Supplementary Table 6**).

Consider a set *s* containing *m* generated UMIs. A newly generated UMI *u* needs to undergo verification to ascertain that it differs by at least a Hamming distance of *d* from any existing UMI within set *s*.

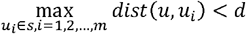

#### Cell barcodes

Cell barcodes are instrumental in pinpointing the precise cell of origin for a molecule, and they can be identified through reference to a whitelist. As an alternative to a library of synthesized barcodes, 10X Genomics V2 and V3 whitelists can be optionally utilized for generating barcodes.

#### Customized components

Tresor enables the generation of a limitless array of customized components, each with sequences of any desired length and nucleotides drawn at random. Integrating customized components into a sequencing library may be adopted to facilitate validation of their bespoke sequencing experiments.

### Approach for simulation of PCR amplification

Simulation of PCR amplification is often involved with multiple steps in read copying and error generation. At stage 1 for read copying, reads will potentially be subjected to shuffling, sampling (read number sampling and read identifier sampling), and concatenation. At stage 2, error generation is likely to involve error position sampling, error nucleotide sampling, error assignment, error table building, and PCR tree construction. For example, to remove stochastic biases towards the selectivity of a specific bunch of reads, we shuffle reads after each PCR amplification cycle. To efficiently manage many such steps, we employed aspect-orientated programming techniques to claim whether each of the above step is specifically required for different read simulation methods. As listed in **Supplementary Fig. 1**, PCR-tree-based methods undertake the minimum steps, making it much faster during PCR amplification compared with read table-based and error table-based methods. The output will be delivered to another aspect-orientated programming module for simulating sequencing.

### Flexible arrangement and combination between components

Innovations in sequencing rely heavily on numerous experimental trials. To streamline laboratory efforts, conducting computational simulations of early-stage concepts before experimental trials are crucial to provide valuable insights and preliminary data for reference. At this stage, tailored sequencing experiments may be organized, potentially necessitating the creation of new sequencing libraries. To address this, Tresor incorporates a module enabling the flexible design of sequencing libraries, where existing or new components of reads can be arranged and combined as needed. This stands out as Tresor’s most distinctive feature.

### Parameter settings for read simulation

To evaluate the runtime of simulating reads by our proposed framework, we arranged a series of *in silico* experiments at different application scenarios, including PCR cycles, PCR amplification efficiencies, PCR sequencing errors, sequencing errors, and sequencing depth. In scenario, we varied on parameter in a certain range but kept the rest of the parameters fixed. The fixed parameters include PCR cycle (12), amplification efficiency (0.85), PCR error rate (0.0001), sequencing error rate (0.01), sequencing depth (500), permutation times (10), Hamming distance threshold for UMIs (3), number of initial molecules (50), and UMI length (10). The varied parameters include PCR cycle (6-24), amplification efficiency (0.1, 0.2, 0.3, 0.4, 0.5, 0.6, 0.7, 0.8, 0.9, 1), PCR error rate (0.00001, 0.000025, 0.00005, 0.000075, 0.0001, 0.00025, 0.0005, 0.00075, 0.001, 0.0025, 0.005, 0.0075, 0.01, 0.025, 0.05, 0.075, 0.1), sequencing error rate (0.00001, 0.000025, 0.00005, 0.000075, 0.0001, 0.00025, 0.0005, 0.00075, 0.001, 0.0025, 0.005, 0.0075, 0.01, 0.025, 0.05, 0.075, 0.1), sequencing depth (100, 200, 500, 600, 800, 1000, 2000, 3000, 5000). The number of errors was drawn from negative binomial distributions.

## Conclusion

In this work, we have developed a high-performance read simulation tool, Tresor, which aims to computationally bolster preliminary verification of sequencing-related biological discoveries through simulated reads. To satisfy this need, Tresor simulates sequencing reads at multiple application scenarios and sequencing levels. Simulation of PCR amplification has faced a grand challenge in data volume and time complexity. To ensure the optimal speed for simulation, we constructed a multitude of methods and systematically compared their runtime under different conditions. We show that PCR tree-based methods can simulate reads at the fastest speed in all situations. One of the most important features of Tresor is the self-custom read structure, allowing flexible design of sequencing libraries. This is particularly useful to dig up valuable information from sequencing data of the libraries, facilitating innovations in sequencing technology. The read simulation process in Tresor comprises library preparation, PCR amplification, and sequencing, where substitution, insertion, and/or deletion errors can be added.

Our results demonstrate that by constructing a PCR tree, reads can be simulated with a greatly reduced amount of computing power, enabling that computational resources are not necessarily required to allocate for a substantial number of reads that do not remain for sequencing. This allows for amplifying reads at high PCR cycles and thus may assist in the design of sequencing experiments demanding deep PCR cycles, e.g., single-cell sequencing. In addition, we showcase that in our simulated data, errors introduced during PCR amplification can be obviated by the UMI deduplication method (Marx 2017), which makes it consistent with established research findings.

## Data and code availability

The standalone package of Tresor is made available at https://github.com/2003100127/tresor. The data used in this paper can be accessed through https://2003100127.github.io/tresor.

## Author contribution

J.S. conceived this study. J.S. implemented the algorithm, performed the analysis, developed Tresor, and wrote the manuscript. J.S. and A.P.C. revised the paper and acquired funding. All authors have given approval to the final version of the manuscript.

## Acknowledgement

This work was financially supported by the Medical Research Council (MRC) career development fellowship (MR/V010182/1).

