## Supplementary Information for "Tresor: An integrated platform for simulating transcriptomic reads with realistic PCR error representation across various RNA sequencing technologies"

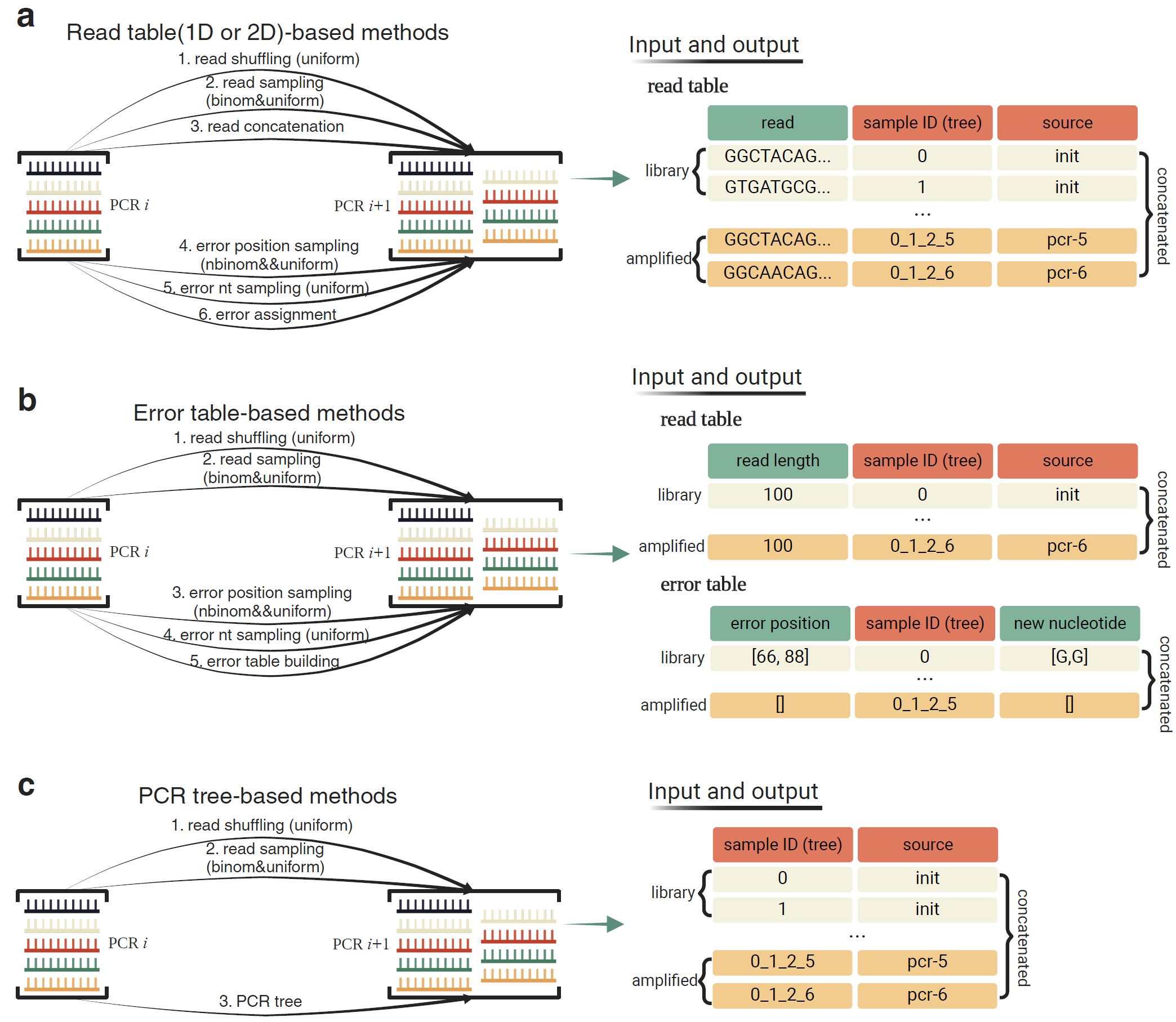

**Supplementary Figure 1**. Overview of read amplification and error generation between every two adjacent PCR cycles using read table-based, error table-based, and PCR tree-based methods.

**Supplementary Table 1.** The pseudocode of the *bfTree* algorithm.

Algorithm 1: *bfTree* - Pseudocode

Initialization: Read dictionary *rd* that maps from read identifiers to read sequences. Function $\boldsymbol{\Phi}$ that samples *n* reads from *rd*. Function $\boldsymbol{\omega}$ that maps from molecule identifiers to all their amplified reads represented by 2D arrays, resulting in PCR trees. Within a 2D array, each vector consists of a series of PCR cycles (e.g., [2, 5, 6]) through which a read is amplified from an original read. The maximum PCR cycle is $\boldsymbol{z}$. Function $\mathcal{g}$ that expands the dimension of each tree vector to $\boldsymbol{z}$ and pads expanded positions by 0 wherever possible. *k* stands for the *kth* original molecule and *at* stands for all its related PCR trees*.* Function $\mathcal{s}$ that simulates substitution errors with a DNA polymerase error rate of $\boldsymbol{\alpha}$. Function $\mathcal{i}$ that simulates insertion errors with an error rate of $\boldsymbol{\beta}$. Function $\mathcal{d}$ that simulates deletion errors with an error rate of $\boldsymbol{\gamma.}$ *repli* stands for a list of indices of an element of interest in a given list.

Output: A list of UMIs *rl*.

Procedure

*rd* ← read dictionary

*rs* ← $\boldsymbol{\Phi}$(*rd*, *n*)

*rsd* ← $\boldsymbol{\omega}$(*rs*, *n*) # PCR tree construction

*rl* ← 2D empty list

bfTAB ← 2D empty list

rcTAB ← 2D empty list

FOR *k*, *at* in *rsd* do

*r* ←*rd*[*k*]

*cache* ← {}

FOR *tree* in *at* do

cc ← max(length(tree), $\boldsymbol{z}$)

tree ← $\mathcal{g}$(*tree*)

update *tree* in *at*

END FOR

\# Boolean table construction

FOR *j* in *at.shape[1]* do

*reple* ← FindDuplicates(*at[:, j]*)

FOR *i* in *at[:,j]* do

IF *i* in *reple* then

*repli* ← LocaliseRepeatIndices(*at[i, j], at[:, j]*)

IF *at[i,j-1] is* False then

bfTAB*[i,j]* ← False

ELSE

IF $\boldsymbol{\forall x\notin}$ *at[ii,j-1]* $\boldsymbol{ii\neq i}$, if $\boldsymbol{x}$=*at[i,j-1],* bfTAB*[ii,j]* is True then

bfTAB*[i,j]* ← True

ELSE

bfTAB*[i,j]* ← False

END IF

END IF

ELSE

bfTAB*[i,j]* ← False

END IF

END FOR

END FOR

\# tracing reads and assigning errors

FOR *j* in *at.shape[1]* do

*cache* ← {}

FOR *i* in *at[:,j]* do

IF bfTAB*[i,j]* is True then

IF *i*$\boldsymbol{\notin}$*cache keys* then

IF $\boldsymbol{\alpha}$ exists then

*r* = $\mathcal{s}$(*r*, $\boldsymbol{\alpha}$)

IF $\boldsymbol{\beta}$exists then

*r* = $\mathcal{i}$(*r*, $\boldsymbol{\beta}$)

IF $\boldsymbol{\gamma}$ exists then

*r* = $\mathcal{d}$(*r*, $\boldsymbol{\gamma}$)

cache[*i*] ← *r*

append *r* to *rl*

ELSE

append cache[*i*] to *rl*

END IF

ELSE

IF $\boldsymbol{\alpha}$ exists then

*r* = $\mathcal{s}$(*r*, $\boldsymbol{\alpha}$)

IF $\boldsymbol{\beta}$exists then

*r* = $\mathcal{i}$(*r*, $\boldsymbol{\beta}$)

IF $\boldsymbol{\gamma}$ exists then

*r* = $\mathcal{d}$(*r*, $\boldsymbol{\gamma}$)

append *r* to *rl*

END IF

END FOR

END FOR

END FOR

End Procedure

**Supplementary Table 2.** The pseudocode of the *spTree* algorithm.

Algorithm 2: *spTree* - Pseudocode

Initialization: Read dictionary *rd* that maps from read identifiers to read sequences. Function $\boldsymbol{\Phi}$ that samples *n* reads from *rd*. Function $\boldsymbol{\varphi}$ to map from molecule identifiers to all their amplified reads represented by PCR trees (e.g., 2_5_6). *k* stands for the *kth* original molecule and *st* stands for all its related PCR trees*.* Function $\mathcal{s}$ that simulates substitution errors with a DNA polymerase error rate of $\boldsymbol{\alpha}$. Function $\mathcal{i}$ that simulates insertion errors with an error rate of $\boldsymbol{\beta}$. Function $\mathcal{d}$ that simulates deletion errors with an error rate of $\boldsymbol{\gamma.}$

Output: A list of UMIs *rl*.

Procedure

*rd* ← read dictionary

*rs* ← $\boldsymbol{\Phi}$(*rd*, *n*)

*rsd* ← $\boldsymbol{\varphi}$(*rs*, *n*)

*rl* ← empty list

FOR *k*, *st* in *rsd* do

*r* ←*rd*[*k*]

*cache* ← {}

FOR *tree* in *st* do

split *tree* according to "_"

*C* ← *k*

FOR *pcr* in *tree* do

IF *pcr* exists then

*C* ← *C* + '_' + *pcr*

ELSE

keep *C* unmodified

END IF

IF *C* in *cache* then

*r* = *cache*[k_]

ELSE

IF $\boldsymbol{\alpha}$ exists then

*r* = $\mathcal{s}$(*r*, $\boldsymbol{\alpha}$)

IF $\boldsymbol{\beta}$exists then

*r* = $\mathcal{i}$(*r*, $\boldsymbol{\beta}$)

IF $\boldsymbol{\gamma}$ exists then

*r* = $\mathcal{d}$(*r*, $\boldsymbol{\gamma}$)

END IF

append *r* to *rl*

END FOR

END FOR

END FOR

End Procedure

**Supplementary Table 3.** The pseudocode of the *ReadPer* algorithm

Algorithm 3: *ReadPer* - Pseudocode

Initialization: Read dictionary *rd* that maps from read identifiers to read sequences. Function $\mathcal{s}$ that simulates substitution errors with a DNA polymerase error rate of $\boldsymbol{\alpha}$. Function $\mathcal{i}$ that simulates insertion errors with an error rate of $\boldsymbol{\beta}$. Function $\mathcal{d}$ that simulates deletion errors with an error rate of $\boldsymbol{\gamma.}$

Output: A list of UMIs *rl*.

Procedure

*rd* ← read dictionary

*rl* ← empty list

FOR *r* in *rs* do

*r* ← *rd[kt]*

IF $\boldsymbol{\alpha}$ exists then

*r* = $\mathcal{s}$(*r*, $\boldsymbol{\alpha}$)

IF $\boldsymbol{\beta}$exists then

*r* = $\mathcal{i}$(*r*, $\boldsymbol{\beta}$)

IF $\boldsymbol{\gamma}$ exists then

*r* = $\mathcal{d}$(*r*, $\boldsymbol{\gamma}$)

append *r* to *rl*

END FOR

End Procedure

**Supplementary Table 4.** The pseudocode of the *ReadWhole* algorithm

Algorithm 4: *ReadWhole* - Pseudocode

Initialization: Read dictionary *rd* that maps from read identifiers to read sequences. Function $\mathcal{m}$ that concatenates all reads as a whole and return sequence positions (*pid*) and their related read identifiers (*rid*). Function $\mathcal{s}$ that simulates substitution errors with a DNA polymerase error rate of $\boldsymbol{\alpha}$. Function $\mathcal{i}$ that simulates insertion errors with an error rate of $\boldsymbol{\beta}$. Function $\mathcal{d}$ that simulates deletion errors with an error rate of $\boldsymbol{\gamma.}$

Output: A list of UMIs *rl*.

Procedure

*rd* ← read dictionary

*rl* ← empty list

*rid, pid* ← $\mathcal{m}$(*rd*)

IF $\boldsymbol{\alpha}$ exists then

*rd* = $\mathcal{s}$(*rd, rid, pid*, $\boldsymbol{\alpha}$)

IF $\boldsymbol{\beta}$exists then

*rd* = $\mathcal{i}$(*rd, rid, pid*, $\boldsymbol{\beta}$)

IF $\boldsymbol{\gamma}$ exists then

*rd* = $\mathcal{d}$(*rd, rid, pid*, $\boldsymbol{\gamma}$)

*rl* ← *rd*

END FOR

End Procedure

**Supplementary Table 5.** The pseudocode of the *ErrorTable-s or ErrorTable-m* algorithm

Algorithm 5: *ErrorTable-s* or *ErrorTable-m* - Pseudocode

Initialization: Read dictionary *rd* that maps from read identifiers to read sequences. Position dictionary *pd* and base dictionary *bd*. Error table *et*. *kts* stands for a list containing a string of read identifier and its PCR cycles (e.g., 1_2_5_6, 1 represents the read identifier and 2_5_6 reflects PCR cycle information)*.* Function $\mathcal{s}$ that simulates substitution errors with a DNA polymerase error rate of $\boldsymbol{\alpha}$. Function $\mathcal{i}$ that simulates insertion errors with an error rate of $\boldsymbol{\beta}$. Function $\mathcal{d}$ that simulates deletion errors with an error rate of $\boldsymbol{\gamma.}$

Output: A list of UMIs *rl*.

Procedure

*rd* ← read dictionary

*rl* ← empty list

extract *pd* from *et*

extract *bd* from *et*

FOR *kt* in *kts* do

*r* ← *rd[kt]*

extract PCR cycle information *id* from *kt*

extract read identifier *k* from *kt*

IF $\boldsymbol{\alpha}$ exists then

*r* = $\mathcal{s}$(*r*, $\boldsymbol{\alpha}$)

IF $\boldsymbol{\beta}$exists then

*r* = $\mathcal{i}$(*r*, $\boldsymbol{\beta}$)

IF $\boldsymbol{\gamma}$ exists then

*r* = $\mathcal{d}$(*r*, $\boldsymbol{\gamma}$)

append *r* to *rl*

END FOR

End Procedure

**Supplementary Table 6.** The pseudocode of the algorithm for simulating UMIs.

Algorithm 6: Simulation of UMIs - Pseudocode

Initialization: Similarity threshold *t*. Read number *N*. Hamming distance function *Hamming.*

Output: A set of UMIs *pool*.

Procedure:

*pool* ← empty list

*c* ← 0

FOR *i* ← 0 to *N* do

*flag* ← False

WHILE not *flag*

generate a seed *s* using *c* for UMI *u*

apply a uniform distribution to generate *u*

IF *i* ← 0 then

append *u* to *pool*

ELSE

*distance* ← empty list

FOR *v* in *pool* do

*dist* ← *Hamming*(*u*, *v*)

append *dist* to *distance*

END FOR

IF none of elements in *distance* < *t* then

append *u* to *pool*

*flag* ← True

write *s* to a file

write *u* to a file

ELSE

*c* ← *c* + 1

END ELSE

END ELSE

END WHILE

END FOR

End Procedure
